# RangeShiftR: an R package for individual-based simulation of spatial eco-evolutionary dynamics and species’ responses to environmental change

**DOI:** 10.1101/2020.11.17.384545

**Authors:** Anne-Kathleen Malchow, Greta Bocedi, Stephen C. F. Palmer, Justin M. J. Travis, Damaris Zurell

## Abstract

1. Reliably modelling the demographic and distributional responses of a species to environmental changes can be crucial for successful conservation and management planning. Process-based models have the potential to achieve this goal, but so far they remain underused for predictions of species’ distributions. Individual-based models offer the additional capability to model inter-individual variation and evolutionary dynamics and thus capture adaptive responses.
2. We present RangeShiftR, an R package that provides flexible and fast simulations of spatial eco-evolutionary dynamics and species’ responses to environmental changes. It implements the individual-based simulation software RangeShifter for the widely used statistical programming platform R. The package features additional auxiliary functions to support model specification and analysis of results. We provide an outline of the package’s functionality, describe the underlying model structure with its main components and present a short example.
3. RangeShiftR offers substantial model complexity, especially for the demographic and dispersal processes. It comes with comprehensive documentation and elaborate tutorials to provide a low entry level. Thanks to the implementation of the core code in C++, the computations are fast. The complete source code is published under a public licence, making adaptations and contributions feasible.
4. The RangeShiftR package facilitates the application of individual-based and mechanistic modelling to eco-evolutionary questions by operating a flexible and powerful simulation model from R. It allows effortless interoperation with existing packages to create streamlined workflows that can include data preparation, integrated model specification, and results analysis. Moreover, the implementation in R strengthens the potential for coupling RangeShiftR with other models.

## Introduction

Under anthropogenic exploitation and rapid environmental changes, one of the most urgent challenges biologists face today is to understand and predict if and how species will persist, by adapting or undergoing changes in their geographic range (Brondizio et al., 2019; McGill et al., 2015). To infer a species’ niche from data and make predictions in space and time, correlative species distribution models (SDMs) are commonly used tools (Elith & Leathwick, 2009; Guisan & Zimmermann, 2000; Qiao et al., 2015). The widespread use of SDMs has been facilitated by accessible and ready-to-use software, most notably Maxent (Phillips et al., 2017) and dedicated R packages such as biomod2 (Thuiller et al., 2009) and dismo (Hijmans et al., 2017). However, these methods often incorporate little ecological theory (Austin, 2007; Guisan & Thuiller, 2005) and usually require making assumptions that are routinely violated in natural observed systems (Elith et al., 2010; Jarnevich et al., 2015; Martínez-Minaya et al., 2018). For example, SDMs assume that species are at equilibrium with their environment and ignore any transient dynamics (Zurell et al., 2016). An alternative that avoids some of these drawbacks is the development and application of mechanistic models, which aim to simulate relevant eco-evolutionary processes such as dispersal, demography and evolution (Cabral et al., 2017; Urban et al., 2016). Despite repeated calls for more mechanistic understanding of range dynamics (Connolly et al., 2017; Kearney & Porter, 2009; Schurr et al., 2012), such models remain underused, arguably due to challenges such as poor availability of the data needed for parametrisation and restricted accessibility of the required software (Briscoe et al., 2019; Dormann et al., 2012).

The ambition for a more prominent representation of process-based models in ecological research led to the development of RangeShifter (Bocedi et al., 2014), an implementation of a flexible individual-based model (IBM) which simulates eco-evolutionary dynamics in a spatially explicit way. It models population dynamics, dispersal, and evolution as interacting processes, organised within a modular structure in which each process has a number of modelling options. This makes RangeShifter a highly adaptable platform with a wide range of applications, including conducting population viability or connectivity analyses (Aben et al., 2016; Henry et al., 2017) and assesing the dynamics of genetic variation across complex landscapes. RangeShifter is a Windows application that can be used via a graphical user interface (GUI) or in a batch mode. The new version 2.0 (Bocedi et al., 2020) adds novel features including the option for dynamic landscapes and a completely revised genetics module. Here, we present RangeShiftR, a package that implements the RangeShifter 2.0 simulation in R (R Core Team, 2020), making it a multi-platform software.

With the RangeShiftR package, we take a step towards a more effortless and accessible use of mechanistic individual-based models. It extends the existing suite of R packages for ecological modelling, which comprises software like the spatially explicit population models steps (Visintin et al., 2020) and demoniche (Nenzén et al., 2012), by a complete and flexible IBM with detailed dispersal dynamics. The package augments the RangeShifter platform with functionality to assist in model specification and output visualisation. As part of the R environment, RangeShiftR offers the powerful potential to interoperate with other packages in order to form integrated workflows, drawing on the extensive functionality for data preparation, output analysis, and easy reporting that is available for R. The actual numeric simulation is implemented entirely in C++ and accessed via Rcpp (Eddelbuettel et al., 2011), thus ensuring high computational performance. RangeShiftR is published under the public license GPLv3 and hence is free to use, modify and share. In order to provide easy access for all users, the package includes extensive built-in documentation and comes with elaborate tutorials presented on the accompanying website (https://rangeshifter.github.io/RangeshiftR-tutorials/).

### Package Structure

The RangeShiftR package inherits its model structure from the underlying RangeShifter platform (Fig. 1). The model requires inputs that include information about the study species and the landscape. These are provided in the form of model choices, parameters and raster maps. The simulation itself is based on a regularly gridded landscape and runs over discrete time steps. It models three main, interacting processes: demography, dispersal, and genetics. Various levels of output can be generated during the simulation and used for analyses. The functions and classes comprising the RangeShiftR package (Fig. 2) reflect this conceptual workflow and can be categorised into the following three groups.

**Figure 1:**
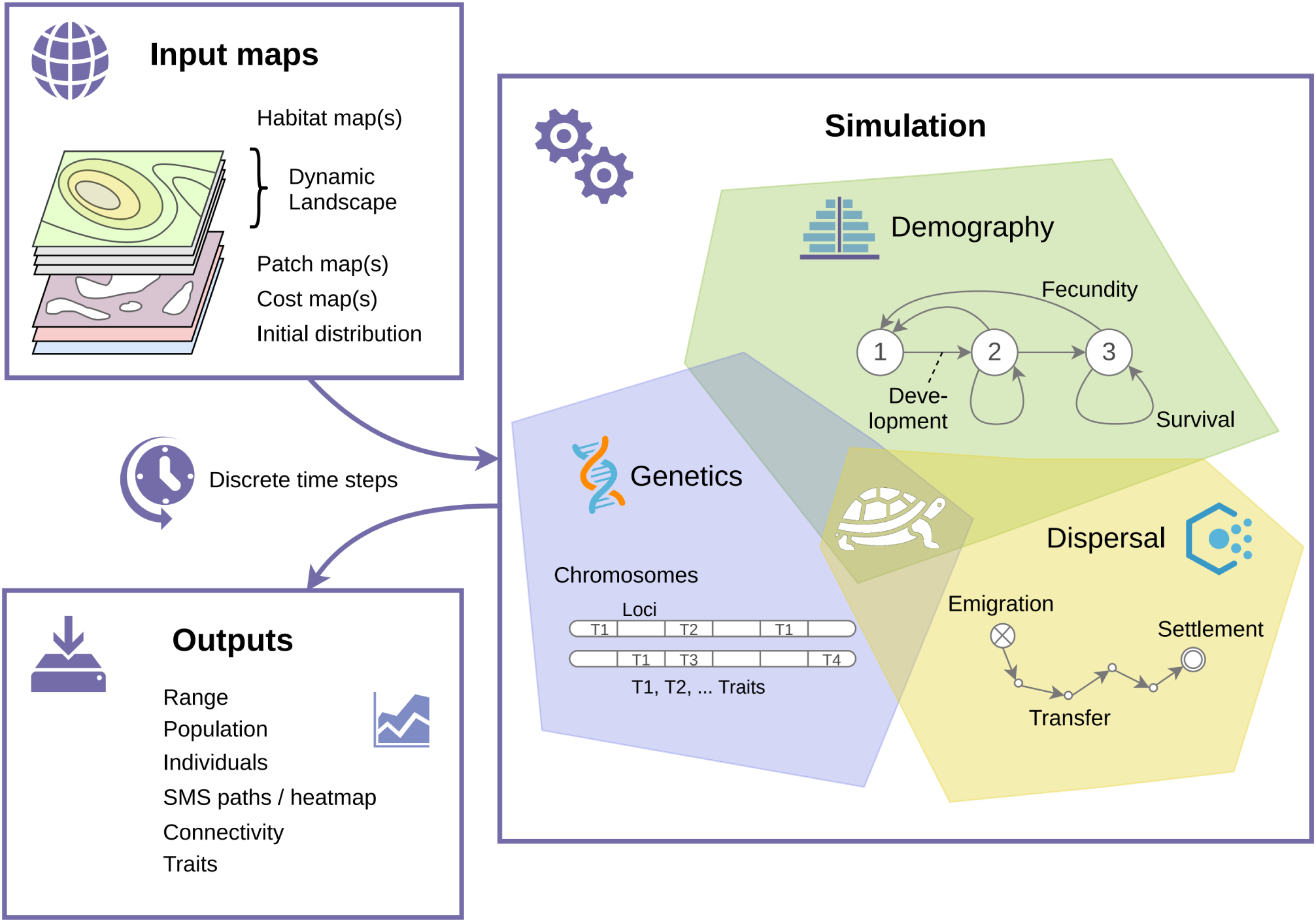
Conceptual overview of a RangeShiftR simulation. The user provides input maps to characterise the landscape, and specifies parameter options that define the three interacting processes of demography, dispersal, and genetics. One option for representing each process is symbolised here as an exemplary model configuration. Different outputs are generated during the simulation and stored in files.

**Figure 2:**
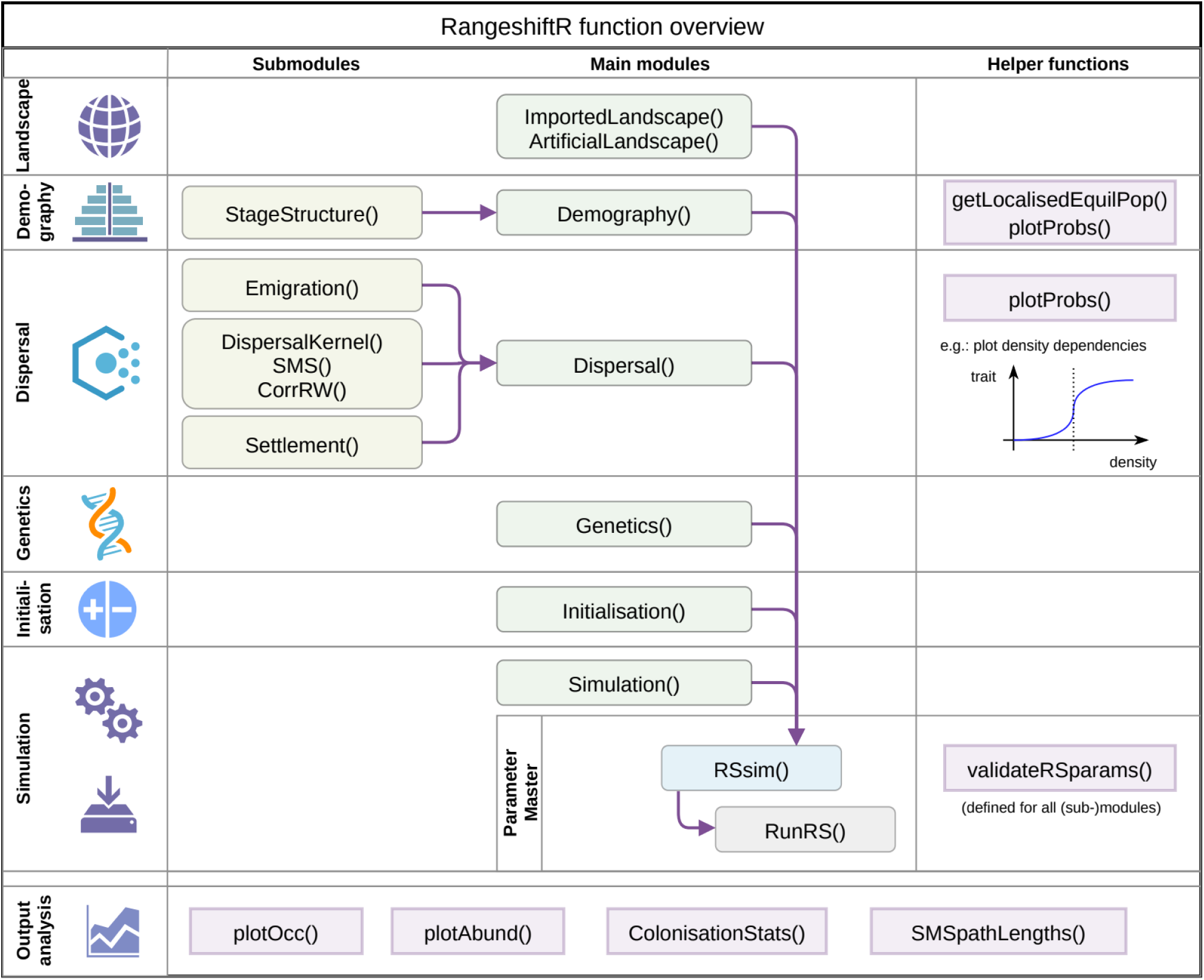
RangeShiftR function overview. The first column introduces the various modules with their respective icons, as reference to Fig. 1. The rounded boxes and arrows in columns 2 and 3 indicate model functions and their respective hierarchical relations. They are class constructors used to define the sub-modules (column 2, yellow) and main modules (column 3, green), which can be combined to a parameter master (blue) to compose the RangeShiftR model. The function RunRS (grey) then starts the simulation. The angled boxes in the last column indicate helper functions that are related to their respective modules. The angled boxes in the bottom row are separate from the columns and itemise the output functions that can be used for processing the simulation results.

### Model functions

A RangeShiftR model is defined by its structure as well as its parameters. The model structure is formed by the assembly of various modules that define the different processes and sub-processes, such as demography or emigration. Each module is represented in R by its own class, whose instances hold the (numeric) values of all model parameters relevant to the respective module. These instances are constructed via the model functions (Fig. 2).

Among the modules exists a number of interdependencies, which are induced by the hierarchy of processes and their sub-processes as well as compatibility restrictions with some options (Bocedi et al., 2014). To reflect this, all modules are organised in a ‘ParameterMaster’ class, that consolidates all components of the model and gives informative error messages or warnings to the user in case of incompatible parameter combinations. An object of this class defines the entire RangeShiftR model and is used for running the simulation.

### Helper functions

Most modules in a RangeShiftR simulation influence each other either directly or indirectly, and the specification of a certain parameter might have implications in various places. Therefore, it can prove challenging to express knowledge about the system by directly specifying numerical parameter values. To aid parameter specification, RangeShiftR includes helper functions to estimate or visualise the effect of some parameters (Fig. 2). For example, they can be used to plot density-dependent demographic or dispersal relationships, whose shape may vary by stage and sex. RangeShiftR also contains the novel functionality to estimate the combined effect of density-dependent population dynamics on a closed population (cf. example below), to guide the choice of adequate levels of demographic rates.

### Output functions

All simulation output is written to text files in the formats provided by RangeShifter. The RangeShiftR package includes dedicated output functions that facilitate the inspection of these results by processing and visualising the output files. This includes, for example, the creation of dispersal heatmaps, which show the number of dispersers that passed through each location, and the computation of statistics such as the occupancy probability and the time to colonisation.

### Simulation Modules

In the modular structure of RangeShiftR, each module represents a different aspect or process of the simulation (Fig. 2), allowing for adaptable levels of model complexity. Below, the main modules are described briefly. For comprehensive documentation, covering all parameters and options, we refer to the package documentation and the RangeShifter manual (Bocedi et al., 2014, 2020).

### Landscape

A RangeShiftR simulation runs on a cartesian grid in which each cell holds information about its cover. This is characterised either by a land type index or by a habitat quality score. The landscape map can be imported from an ASCII raster file or it can be artificially created by a built-in function. Imported landscapes have additional options: they can be patch-based, in which case a second raster file is required to indicate each cell’s patch ID. Additionally, a raster of dispersal resistance values and a presence-absence raster of the initial distribution can be loaded.

### Demography

The modelled demography is determined by two main choices: 1) The population can have overlapping or non-overlapping generations, meaning it can be stage-structured or not. A stage-structure is a sub-module that is represented by its own class and that can be added to the demography module. 2) The population can be modelled using either both sexes or females only. In the former case, various parameters can be sex-specific and the reproductive dynamics may optionally include an explicit mating system.

### Dispersal

The dispersal module has three obligatory sub-modules: ‘Emigration’, ‘Transfer’, and ‘Settlement’, which represent the three explicitly modelled phases of dispersal (Travis et al., 2012). Emigration and settlement can be density-dependent and, if applicable, sex- and stage-specific.

The transfer sub-module offers three options: it can be modelled with a dispersal kernel or with explicit consideration of the movement processes using either the stochastic movement simulator (SMS) (Palmer et al., 2011) or a correlated random walk.

### Genetics

Individuals can carry a genome that they inherit from their parents at birth. The genome may consist of multiple autosomal loci that can either be neutral or coding for traits (Bocedi et al., 2020). Most dispersal parameters can optionally be treated as heritable traits thus allowing evolution of dispersal strategies. The genetic architecture is highly flexible and processes such as recombination, mutation and pleiotropy can be explicitly modelled. Modelling of neutral loci allows explicit and individual-based population genetic simulations to address questions on how environmental features and processes, in interaction with population dynamics and dispersal behaviours, shape the genetic structure and diversity of populations (Manel et al., 2003).

### Initialisation

The initial state of the simulation in the starting year can be defined in three different ways: with an initial distribution map, with a list of individuals and their location, or at a given density in randomly selected locations.

### Simulation

This module specifies the general simulation settings like the number of simulated time steps (years) and replicates, the types of generated output, and some more specialised options, such as imposing a (shifting) gradient or enabling environmental stochasticity.

## Using RangeShiftR

As widely applicable simulation software, RangeShiftR aims to provide easy access via a range of resources to support the user: all functions are comprehensively documented on R help pages, an extensive user manual is available online, and the webpage (https://rangeshifter.github.io) features a support forum as well as a collection of detailed tutorials that illustrate the model’s scope and introduce the available modelling options. The tutorials include adaptations of the three original RangeShifter examples (Bocedi et al., 2014), accompanied by sample code for analysis and visualisation. Additionally, we provide a fourth tutorial that demonstrates novel features of RangeShifter 2.0 (Bocedi et al., 2020) by simulating the range dynamics of a species in a changing landscape. Here, we present its shortened form to introduce the RangeShiftR syntax.

### Landscape

When using the novel feature of dynamic landscapes, we specify the file names of the changing habitat maps, their corresponding patch files, and the order of years in which these become effective. All maps are imported as ASCII rasters by the function ‘ImportedLandscape()’. Further arguments are the number of land cover types ‘Nhabitats’, their respective density-dependence ‘K_or_DensDep’, and the initial distribution map. The value of ‘K_or_DensDep’ depends on the demography: for a non-structured population it is interpreted as carrying capacity (K) whereas for a stage-structured population it represents the strength of demographic density dependence (1/b).

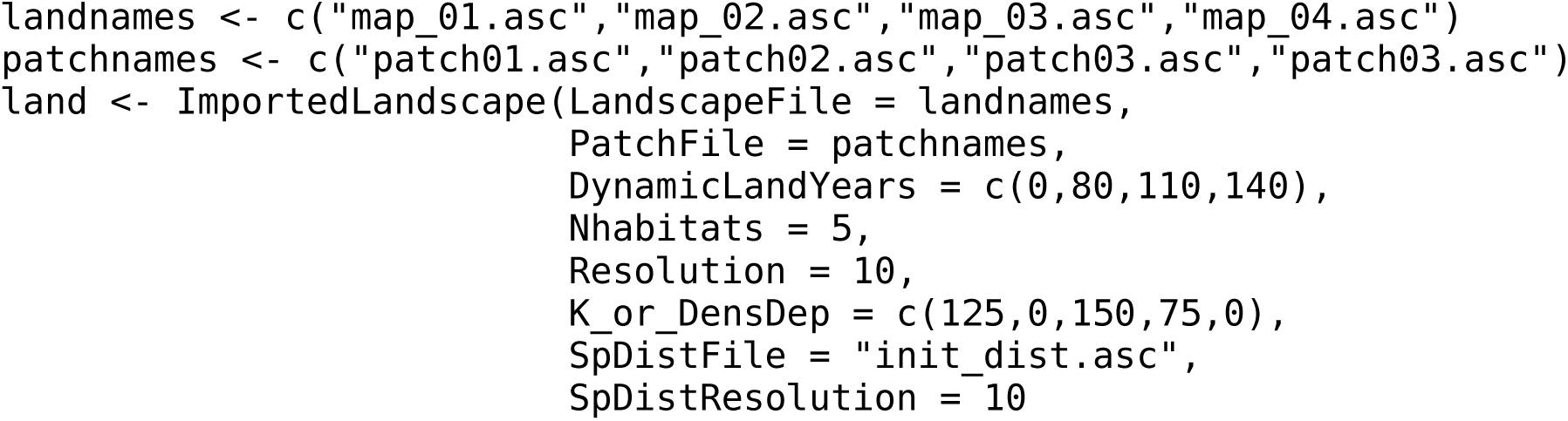

### Demography

The population model is set up to use explicit sexes and a stage-structure, i.e. generations are overlapping. In the ‘Demography()’ module the coded argument ‘ReproductionType’ determines whether both sexes are modelled. The ‘StageStructure()’ sub-module takes the transition matrix and sets optional density-dependencies on the sub-processes of fecundity, survival and development.

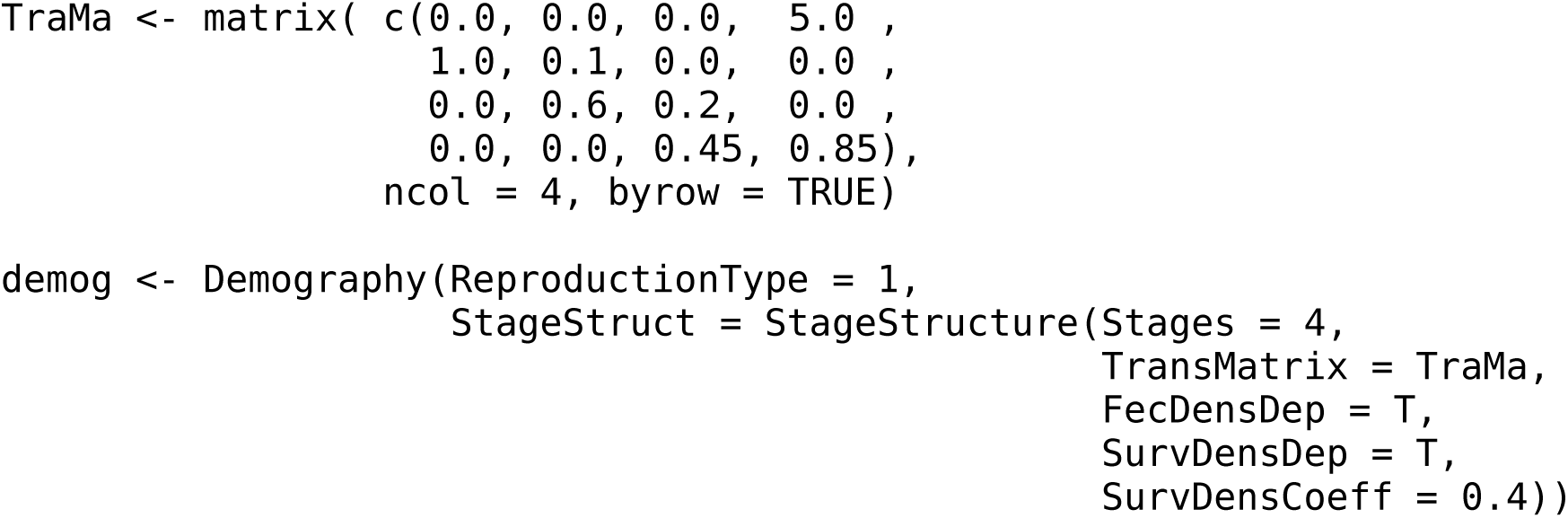

The helper function ‘getLocalisedEquilPop()’ can assist in understanding how the demographic rates of the ‘Demography()’ module and the local density-dependence (1/b) affect the simulated abundances:

~~~
getLocalisedEquilPop(demog = demog, DensDep_values = seq(50,300,50))
~~~

It simulates a time series of the density (in individuals per hectare) of a single closed population for varying values of 1/b (given by ‘DensDep_values’). This is achieved by repeated matrix multiplication with the density-dependent transition matrix until an equilibrium is reached. The function returns these equilibrium densities by stages at the given density-dependence values and generates a bar graph (Fig. 3a). The generated densities approximate the equilibrium densities of a closed patch in the RangeShiftR simulation, and can thus be used to guide the choice of the parameter 1/b. However, the matrix approach neglects stochasticity, the scheduling of survival and reproduction, and the integer units of abundance, so that the quality of the estimate is lower for smaller populations.

**Figure 3:**
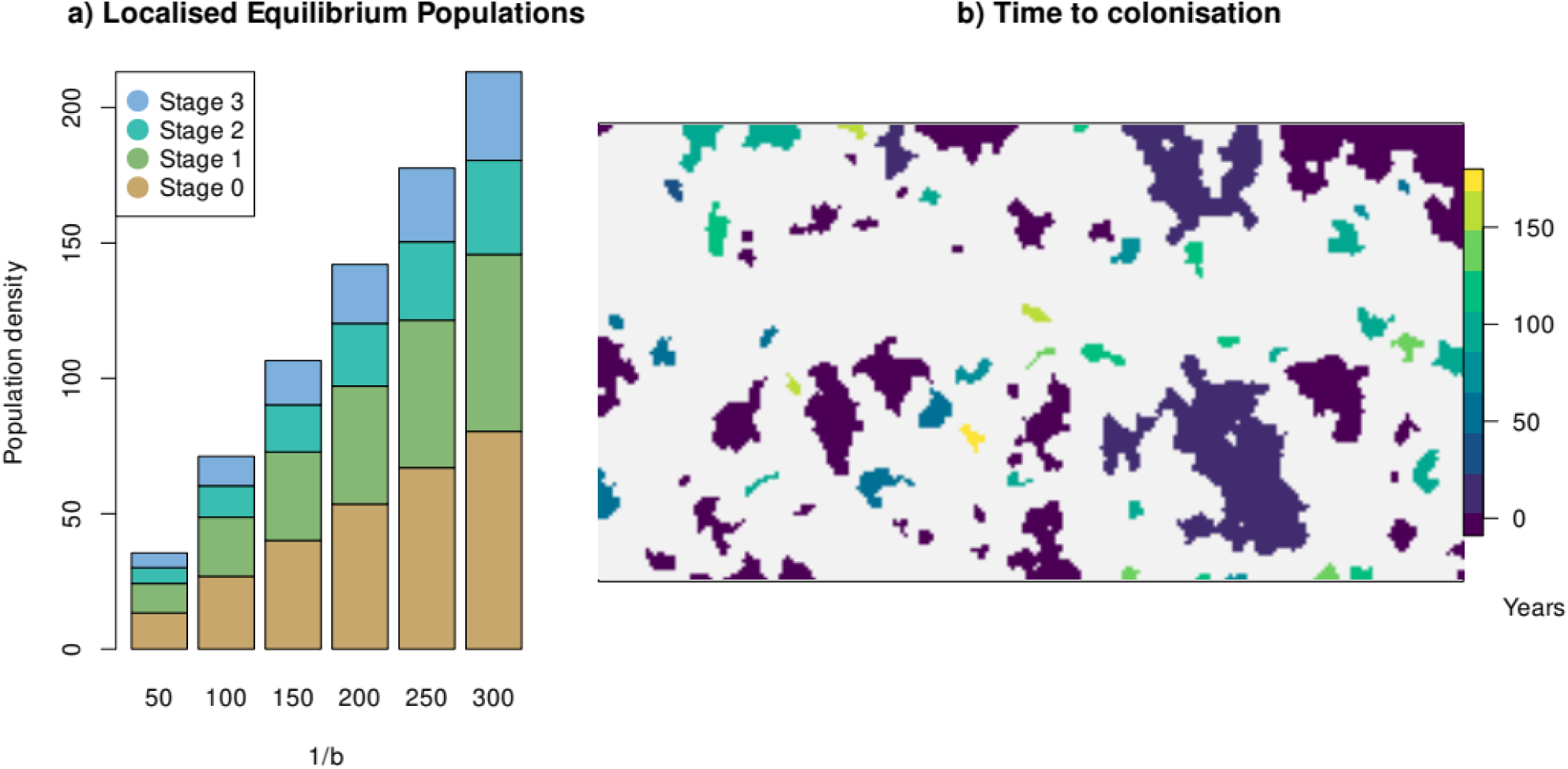
RangeShiftR example. (a) Bar graph generated by the helper function ‘getLocalisedEquilPop()’, showing the approximated equilibrium densities classified by stages over the parameter 1/b (both in units of Inds/ha). They serve as a quick approximation to assess the effect of density-dependent demographic rates. (b) Raster generated by the output function ‘ColonisationStats()’, showing the average time to colonisation.

### Dispersal

The three phases of dispersal are defined as sub-modules before assembling them in the dispersal module. The emigration probability is modelled as stage- and density-dependent, therefore we provide a matrix with one row per stage containing three parameters each, which define how emigration probability relates to population density:

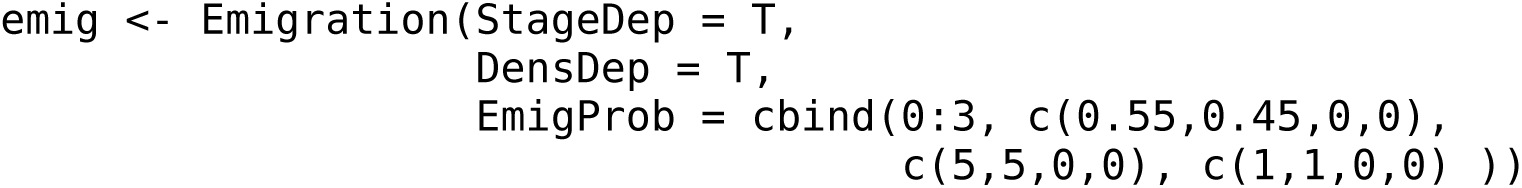

The transition phase uses SMS, setting a dispersal bias (entire second line), habitat-specific dispersal resistances, and a constant per-step mortality:

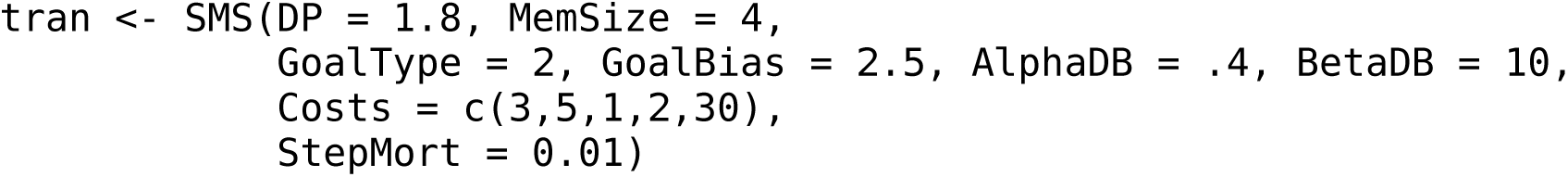

The settlement module defines the minimum and maximum number of steps permitted and sets the mate-finding requirement:

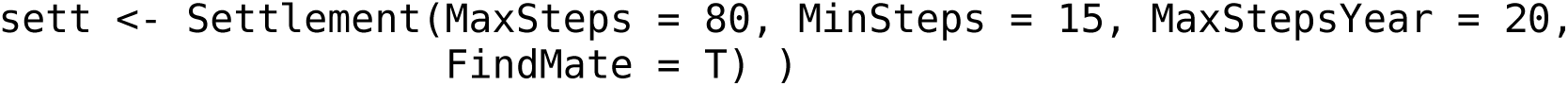

Now, the previously defined sub-modules can be combined in the ‘Dispersal()’ module:

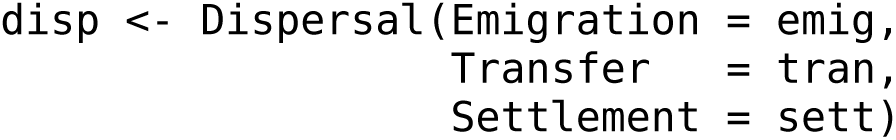

### Genetics

The genetics module is optional and we leave it disabled here (but see Bocedi et al. (2020) for an example of this functionality). Although this implies missing inter-individual variation in dispersal traits, individuals are still characterised by their sex, stage and age.

### Initialisation

The simulation is initialised in all locations indicated by the initial distribution map (in the landscape module) at a given density. Further, the stage- and age-distributions of the initial population are set.

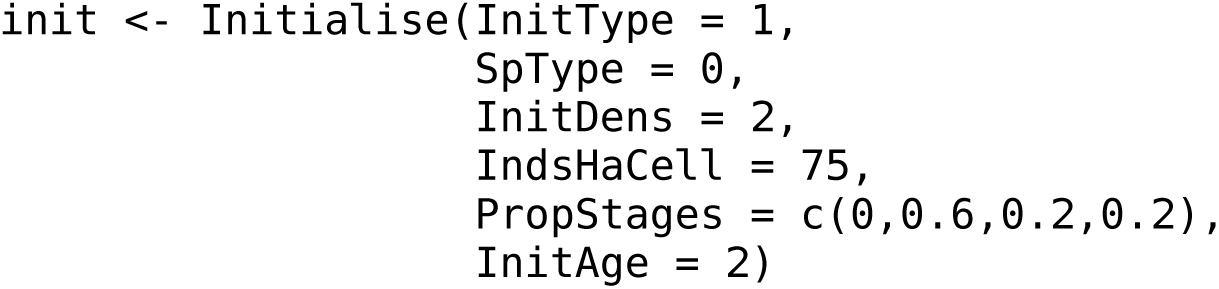

### Simulation

The simulation runs for 200 years and over twenty replicates. The population, range and SMS paths outputs are enabled and will be generated at the given time intervals:

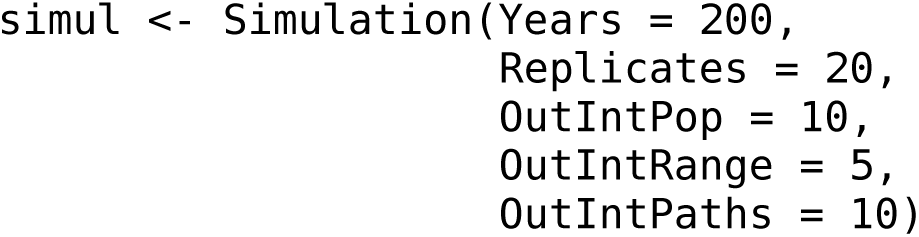

### Model run and results

All defined model components are combined into the parameter master with ‘RSsim()’. Every RangeShiftR simulation is defined by an instance of this class and the path to its directory and is run using ‘RunRS()’:

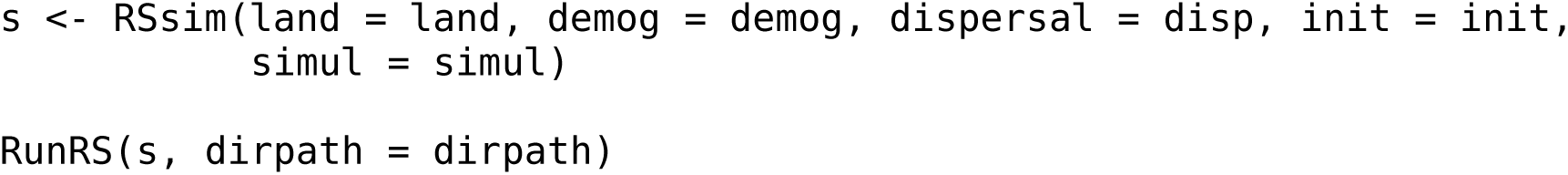

The simulation output is written to text files in the ‘Outputs’ folder of the directory, which can be visualised and processed using the auxiliary output functions. For example, Fig. 3b shows the result of the following function that calculates, among other things, the time to colonisation and maps it onto the landscape:

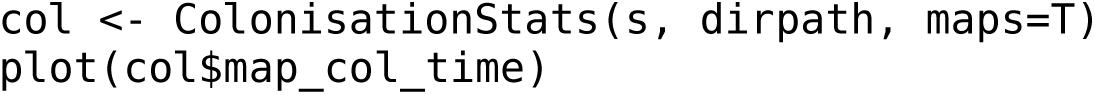

In the resulting plot, the non-suitable landscape matrix appears grey and all habitat patches are coloured according to their averaged time to colonisation over all replicates. In this example, smaller patches tend to get colonised later than larger ones.

## Discussion

RangeShiftR provides, for the first time, an open-source eco-evolutionary numerical simulation platform that can be controlled and analysed entirely from R. Building on the established RangeShifter platform (Bocedi et al., 2014, 2020), it offers a high degree of model complexity, especially for the demographic and dispersal processes. Despite this complexity, straightforward use of the software is provided through the provision of helper functions and comprehensive documentation.

The RangeShifter GUI and the RangeShiftR package constitute two complementary entities, as they represent alternative interfaces to the same software core. The GUI version offers an intuitive handling of the model and visual tracking of simulation outcomes, making it particularly suited for the use by stakeholders or for undergraduate education. The RangeShiftR package, on the other hand, is especially useful for research purposes. It offers transparent, reproducible workflows, as the entire simulation can be scripted in R, along with the visualisation and post-analysis of simulation results. This also facilitates large-scale parameter comparisons, as in sensitivity and robustness analyses. The use of Rcpp (Eddelbuettel et al., 2011) allows running of the simulation in a C++ module and thereby yields high performance, while the integration in R makes RangeShiftR available for multiple platforms and provides the infrastructure for parallel and cluster computing without having to adapt the C++ backend.

RangeShiftR holds many opportunities for interoperation with other R packages. Firstly, it can be readily integrated with packages for describing the landscape context (e.g. raster; (Hijmans & van Etten, 2016)) or species distribution modelling (e.g. biomod2; (Thuiller et al., 2009), sdm; (Naimi & Araújo, 2016)). Secondly, it permits coupling of different model types, as exemplified by coupling RangeShifter with the land-use model CRAFTY (Murray-Rust et al., 2014; Synes et al., 2019).

Thirdly, it enables integrated use with existing methodological devices, like inverse parameterisation through Bayesian inference, for example using the package BayesianTools (Hartig et al., 2017) RangeShiftR complements the existing toolbox of R packages for ecological simulations by a powerful individual-based eco-evolutionary modelling platform. It offers some important features that have not been available so far. Existing R implementations of spatially-explicit population modelling frameworks, such as the recently published package steps (Visintin et al., 2020) or the demoniche package (Nenzén et al., 2012), are population-based. By contrast, RangeShiftR is individual-based and hence allows for an explicit representation of genetics and evolutionary dynamics. The package vortexR (Pacioni & Mayer, 2017) implements post-analysis functions for the prominent Vortex model (Lacy, 1993) that is also individual-based and commonly applied for population viability analysis (PVA). Here, RangeShiftR provides a useful alternative that allows conducting spatially-explicit PVA under more complex dispersal assumptions.

RangeShiftR can help overcome some of the challenges that have prevented more widespread use of mechanistic range models (Briscoe et al., 2019) by offering high accessibility. In the future, we plan to enhance the platform further to improve forecasts under global change. For example, RangeShiftR is currently restricted to modelling a single species only and does not incorporate species interactions. Moreover, the model operates on a single habitat layer that contains either land types or habitat quality. Therefore, demographic rates are related to the environment only indirectly via the user-defined carrying capacities or density dependence coefficients. Lastly, the genetics module is currently restricted to modelling evolution of dispersal traits while demographic traits cannot evolve. Thus, potential future extensions of the platform will involve explicitly modelling species interactions, demography-environment relationships (Pagel & Schurr, 2012) and genetic evolution of demographic traits. As the code is open source, there is now an opportunity for a broad community of researchers and modellers to contribute to representing these important processes in future versions of the platform.

The RangeShiftR package constitutes an important step towards making frameworks for modelling range dynamics under global change accessible to a wider audience (Lurgi et al., 2015; Schurr et al., 2012; Zurell et al., 2016). We hope that this will inspire a more widespread use of mechanistic distribution models, for example to guide conservation efforts and ecosystem management, and facilitate more seamless integration with other modelling tools.

## Acknowledgements

AM and DZ were supported by Deutsche Forschungsgemeinschaft (DFG) under grant agreement No. ZU 361/1-1. GB was supported by the Royal Society University Research Fellowship.

## Authors’ contributions

AM and DZ conceptualised the modular RangeShiftR package design. GB and SP mainly developed the C++ core code. All authors were involved in key decisions taken during the development of the package. AM primarily wrote, documented and published the RangeShiftR package and led the writing of the manuscript. All authors contributed critically to the drafts and gave final approval for publication.

## Data Availability

The entire source code of the RangeShiftR package is publicly available from GitHub and is licenced under the GNU general public license, version 3 (GPLv3): https://github.com/RangeShifter/RangeShiftR-package

